# Upregulation of the NKG2D ligand ULBP2 by JC polyomavirus infection promotes immune recognition by natural killer cells

**DOI:** 10.1101/2023.05.31.543006

**Authors:** Stephanie Jost, Jenny Ahn, Sarah Chen, Taylor Yoder, Kayitare Eunice Gikundiro, Esther Lee, Simon B. Gressens, Kyle Kroll, Melissa Craemer, G. Campbell Kaynor, Michelle Lifton, C. Sabrina Tan

## Abstract

JC polyomavirus (JCPyV) establishes a chronic infection in 70-90% of the world’s population. In immunocompetent individuals, JCPyV chronic infection is asymptomatic and not associated with diseases. However, JCPyV causes progressive multifocal leukoencephalopathy (PML), a potentially fatal complication of severe immune suppression due to monoclonal antibody treatments for cancer, autoimmune diseases and transplantation, or due to uncontrolled HIV infection. There is currently no effective treatment against PML and novel immunotherapies are urgently needed to decrease the morbidity and mortality caused by JCPyV. The risk of developing PML increases with loss of immune control by JCPyV-specific T cells and antibodies. Natural killer (NK) cells play critical roles in defense against viral infections, yet NK cell contribution to the control of JCPyV infection remains largely unexplored. Here, we first compared NK and T cell responses against JCPyV VP1 peptide pools. In about 40% of healthy donors, we detected robust CD107a upregulation and IFN-γ production by NK cells, extending beyond T cell responses. Next, using a novel flow cytometry-based killing assay, we showed that co-culture of NK cells and JCPyV-infected astrocyte-derived SVG-A cells leads to a 60% reduction in infection, on average. Expression of ligands for the activating NK cell receptor NKG2D was modulated in JCPyV-infected cells, with overall enhanced expression of ULBP2. To evaluate the impact of NKG2D triggering on NK cell-mediated elimination of JCPyV-infected cells, we performed co-cultures in the presence of NKG2D blocking antibodies, which resulted in decreased NK cell degranulation. Altogether, these findings suggest NKG2D-mediated activation may play a key role in controlling JCPyV replication and may be a promising immunotherapeutic target to boost NK cell anti-JCPyV activity.

**Author Summary:** The human polyomavirus JC (JCPyV) infects most people for life but only causes disease in persons with a compromised immune system. In particular, JCPyV reactivation in the brain is responsible for the development of progressive multifocal leukoencephalopathy (PML). There is currently no effective treatment for PML, which is often fatal. Natural killer (NK) cells are effector cells of the innate immune system that play critical roles in defense against viral infections, yet their contribution to the control of JCPyV infection remains largely unexplored. The current study shows that NK cells can eliminate cells infected with JCPyV and that immune recognition is partly mediated by NKG2D, an activating ligand expressed on NK cells, and its binding to ULBP2, a stress-induced ligand expressed on infected cells. Our findings provide new insights into immune mechanisms involved in JCPyV immunity, and unveil opportunities to harness NK cell function in future therapeutic strategies to target JCPyV.

## Introduction

The majority of the adult human population worldwide carries the human polyomavirus JC (JCPyV) [1]. Primary JCPyV infection occurs early in life and results in a persistent, lifelong, asymptomatic infection within the kidney tubular epithelial cells, where the virus reproduces benignly with occasional shedding in the urine [2]. JCPyV can spread to secondary sites, including the bone marrow, lymphoid tissues and the brain [3, 4]. The precise processes underlying entry of JCPyV in the brain have not been completely elucidated and proposed mechanisms include brain transmigrated leukocytes that are either infected with JCPyV or carry viral particles at their surface [5]. JCPyV infection usually does not have significant clinical consequences in immunocompetent individuals. However, prolonged and severe immunosuppression or immunomodulation promotes viral reactivation and increases the risk of developing progressive multifocal leukoencephalopathy (PML). PML is a rare but often fatal infection of oligodendrocytes - the myelinating cells of the central nervous system - by JCPyV. While PML is most common among people living with HIV (PLWH) [6, 7], with up to 80% PML patients being HIV-positive [8], there is a growing number of patients with autoimmune diseases, such as people with multiple sclerosis, at risk for PML [5, 9–12].

There is currently no effective treatment against JCPyV and attempts to treat PML with medications previously approved for other diseases, such as mefloquine, have all failed [13–16]. Furthermore, rapidly restoring the immune functions in the central nervous system can also be fatal due to immune reconstitution inflammatory syndrome [17–20]. Thus, there is an urgent need for novel immunotherapeutic interventions to specifically enhance immune control of JCPyV. Impaired immunity is key in the development of PML. In particular, JCPyV-specific CD8+ T cells are crucial in curtailing JCPyV replication to recover from PML and CD4+ T cell counts are associated with prognosis in PML [21–27]. Based on these observations, recent efforts in PML treatment trials have focused on reviving T cell response using T cell checkpoint inhibitor and allogeneic polyomavirus-specific T cells [28] or infusions of cytokines such as IL-7 and IL-15 superagonist [29, 30]. Results of these studies showed potential efficacy in subgroups of patients, indicating other immune factors, in addition to T cells, are necessary to treat PML in all patients. While it is known that risk of PML increases with the loss of immune control by virus-specific T lymphocytes and antibodies [31, 32], the contribution of natural killer (NK) cells to the containment of JCPyV replication has been suggested in a few studies but remains overall largely unexplored [30, 33, 34]. Classically, NK cells are viewed as nonspecific effector cells of the innate immune system that play critical role(s) in defense against viral infections or nascent neoplasms. Unlike other lymphocytes, NK cells lack antigen-specific receptors but lyse target cells following the integration of inhibitory and activating signals. These signals are generated by an arsenal of germline encoded cell surface molecules, commencing effector functions when activating signals overcome inhibitory ones [35]. NKG2D represents one of the most potent activating NK cell receptors that allow NK cells to discriminate between “self” and a variety of pathological cell states, as engagement with one of its ligands is enough to override inhibitory signals [36]. NKG2D ligands (NKG2DL) consist of MHC class I related chain (MIC) A and B and six UL-16 binding proteins (ULBP1-6), which are typically not expressed in healthy tissues but rather upregulated by cellular stress such as viral infection [37–41]. NKG2D-mediated NK cell responses have been found critical in the control of several viral infections [42–44]. The NKG2D/NKG2DL axis also plays a pivotal role in tumor immunosurveillance and therefore immunotherapeutic strategies targeting the NKG2D pathway are currently under investigation [45–48].

A plethora of studies has provided compelling evidence supporting the significant contribution of NK cells to the immune control of major human viral infections such as CMV, influenza virus, hepatitis C virus and HIV [44, 49–54]. Nevertheless, investigations to better define the role played by NK cells in JCPyV infection are currently lacking and are needed to improve our understanding of PML pathogenesis. Moreover, identification of NK cell subpopulations with enhanced or reduced anti-viral activity against JCPyV could provide novel targets for immunotherapeutic strategies to prevent and treat PML. To fill an important gap in knowledge, in this study we measured NK cell responses against JCPyV and investigated the role of NKG2D, a major activating receptor on NK cells and a checkpoint target for cancer immunotherapies, in NK cell-mediated elimination of JCPyV-infected targets.

## Results

### NK cells display robust responses to JCPyV VP1 peptides

Viral infection induces changes in the HLA peptide repertoire and such alterations can directly [55, 56] or indirectly [57–59] modulate NK cell function. Therefore, NK cell responses can be measured upon stimulation of PBMCs with viral peptide pools or inactivated pathogens and have been proposed to vastly depend on IL-2-secreting effector memory T cells [57–59]. However, to our knowledge, NK cell responses to JCPyV peptide pools have never been investigated. Using PBMCs from JCPyV-seropositive healthy donors, we compared NK and T cell responses against 2 pools of overlapping peptides derived from the JCPyV capsid protein VP1 by intracellular cytokine staining (ICS) (S1 Fig, Fig 1). Previous studies reported T cell responses against JCPyV using ICS, however optimal detection of T cell responses often relied on *ex vivo* expansion of JCPyV-specific T cells for 10 to 14 days of culture in the presence of JCPyV peptides and IL-2 [31]. To avoid culture conditions that could significantly alter and misrepresent primary NK cell responses to JCPyV, we performed a 6h assay in the absence of exogenous IL-2. We observed positive responses to VP1 (defined as at least twice above the background) by bulk NK cells from over 40% of donors, with proportions of IFN-γ+ NK cells being overall higher against the peptide pool encompassing the 161-253 and 257-341 regions of VP1. Using this assay, T cell responses could be directly measured without the need for prior amplification of antigen-specific T cells, yet NK cell responses were consistently more potent than T cell responses.

**Fig 1.**
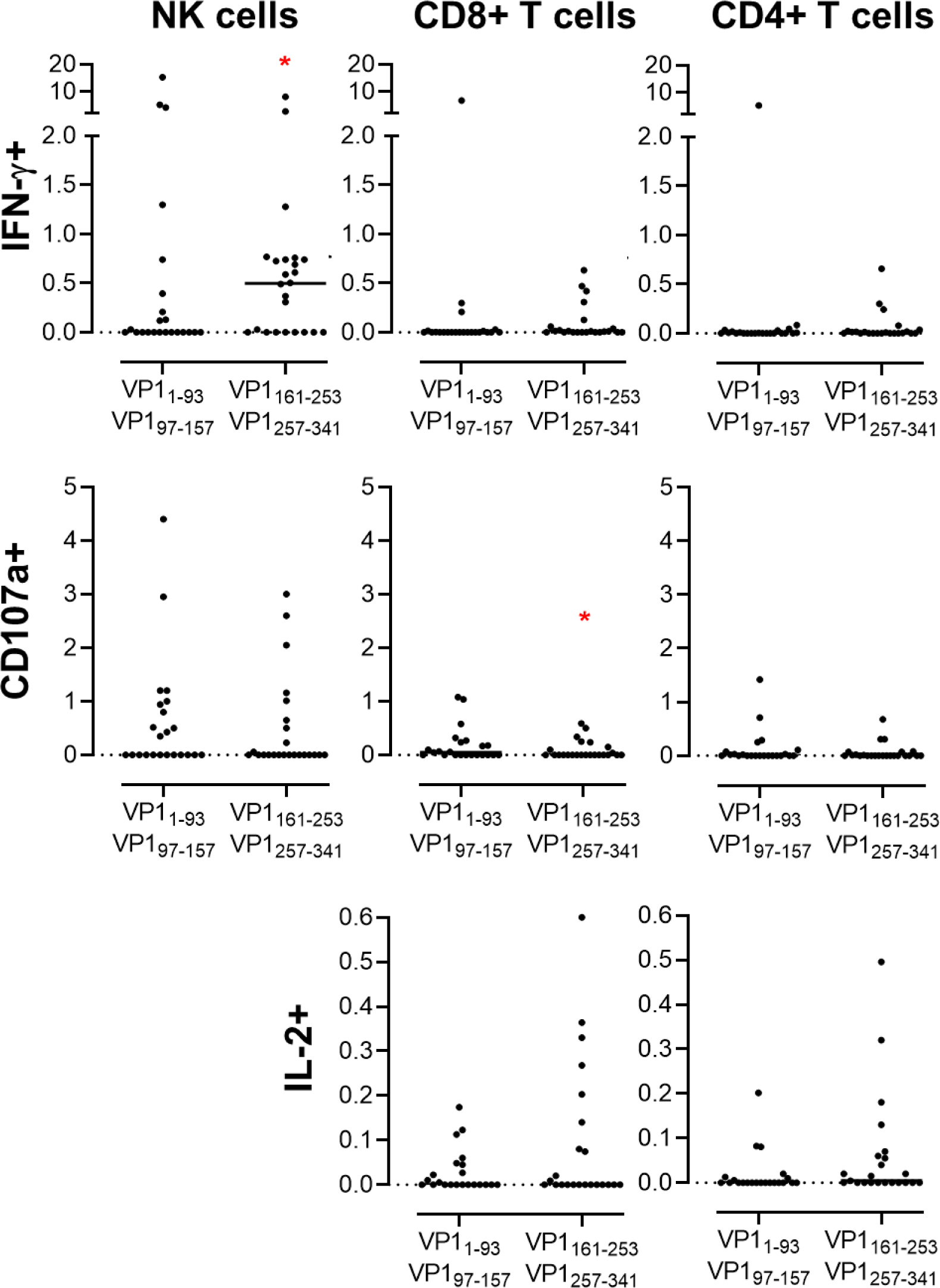
NK and T cell responses to JCPyV VP1 in healthy donors by ICS. PBMC from 24 healthy donors were incubated for 6h with 2µg/mL of pools of peptides spanning JCPyV VP1 in the presence of conjugated anti-CD107a antibodies. BD GolgiStop and GolgiPlug were added for the last 2h of culture. The peptides were divided into two sequential pools as follows: VP1 1-93 + VP1 97-157 (n=48); VP1 161-253 + VP1 257-341 (n=49). Dead cells were excluded using a viability dye. Dot plots represent proportions of CD107a+, IFN-y+ or IL-2+ T or NK cells in response to JCPyV after subtracting proportions of unstimulated cells positive for each marker. Bars represent the median. * p<0.05 compared to unstimulated.

### NK cells efficiently kill JCPyV-infected SVG-A cells

Beyond modulation of NK cell function by changes in the HLA peptide repertoire, recognition of virally infected cells by NK cells mainly rely on downregulation of ligands for inhibitory NK cell receptors (i.e., select HLA class I molecule) and upregulation of ligands for activating receptors (i.e., NKG2DL). To evaluate the overall ability of NK cells to directly recognize and eliminate JCPyV-infected cells, we developed a new flow cytometry-based assay to measure NK cell killing of human fetal astroglial cells immortalized with SV40 (SVG-A) and infected with JCPyV (M1-SVEΔ) (virus originally made in Dr Eugene Major’s lab and provided by Campbell Kaynor, Biogen)[60, 61]. JCPyV M1-SVEΔ is a chimeric polyomavirus derived from the MAD1 strain with replaced regulatory regions from SV40, which display enhanced replication in cell culture [61] (Fig 2A). We show that co-culture of primary NK cells and JCPyV-infected SVG-A cells leads to, on average, a 4-fold decrease (range 1.2-20) in infected cells (Fig 2B) and 60% killing (Fig 2C). While there were inter-individual variations in NK cell-mediated killing of infected SVG-A cells, NK cells from all donors were able to mediate cytotoxic responses against JCPyV. Fifty percent of the donors were seronegative for JCPyV, yet as expected, killing of JCPyV-infected SVG-A cells by NK cells was not influenced by the JCPyV serostatus (data not shown). Of note, CD107a upregulation in NK cells has been shown to correlate with NK cell-mediated cytotoxicity (41, 42) and accordingly, a significant CD107a upregulation was observed in NK cells exposed to JCPyV-infected SVG-A cells (Fig 2D), independently of addition of protein transport inhibitors during the co-culture (S2 Fig). In sum, these data strongly suggest that NK cells significantly contribute to regulating JCPyV replication.

**Fig 2.**
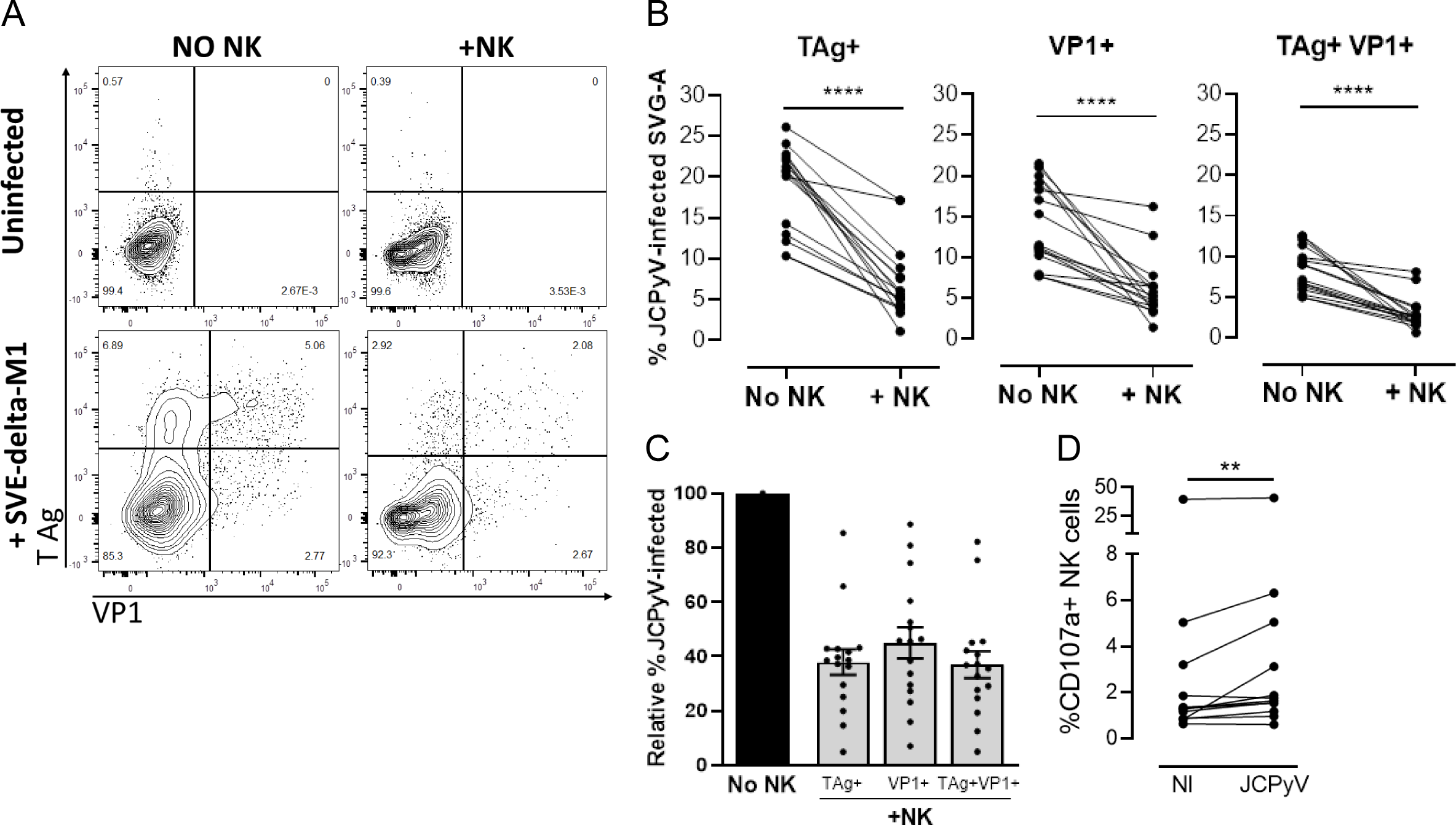
Development of a flow-cytometry-based assay to measure NK cell cytotoxicity against JCPyV-infected cells. SVG-A cells were infected with M1-SVEΔ, which on average leads to 25% cells infected after 9-10 days of culture as assessed by the proportions of SVG-A cells positive for the early protein Large T (TAg) and/or the late capsid protein VP1 by intracellular flow cytometry staining. SVG-A cells were co-cultured with purified NK cells from 16 healthy donors at 10:1 E:T ratio for 6h. (A) Representative flow cytometry plots of uninfected and infected SVG-A cells stained with antibodies targeting TAg and VP1 after gating on live cells in the presence or absence of NK cells. (B) Proportions of TAg+, VP1+ or Tag+VP1+ SVG-A cells after co-culture with NK cells compared to those cultured without NK cells. (C) Relative proportions of JCPyV-infected SVG-A cells in the presence of NK cells. (D) Proportions of CD107a+ NK cells from 13 healthy donors co-cultured with uninfected (NI) SVG-A cells or SVG-A cells infected with M1-SVEΔ (JCPyV) for 6h in the presence of conjugated anti-CD107a antibodies. **p<0.01; **** p<0.0001.

### NKG2D and its ligand ULBP2 play a key role in NK cell responses against JCPyV-infected cells

NKG2D is an activating receptor expressed on most NK cells which is a major regulator of NK cell function [35] and represents a checkpoint target for immunotherapies currently in development [62]. A role of NKG2D in immune responses against polyomaviruses has been previously suggested by studies demonstrating that JCPyV and another member of the family, BKPyV, both express an identical miRNA that target ULBP3 [34]. To evaluate the impact of NKG2D in NK cell-mediated lysis of JCPyV-infected cells in our system, we compared cell surface expression of the different NKG2DL on uninfected and JCPyV-infected SVG-A cells and 293T cells, another cell line permissive for JCPyV infection (Fig 3A-B). SVG-A cells expressed MICA and MICB independently of infection, which are likely responsible for the background response by NK cells against uninfected cells (Fig 2D). We consistently found enhanced binding of an antibody that recognizes ULBP2, ULBP5 and ULBP6 on JCPyV-infected cells, suggesting one or several of these receptors are upregulated upon JCPyV infection. To precisely determine how these receptors are independently modulated by JCPyV infection, we quantified mRNA expression levels in uninfected and infected cells using primers specific for each ULBP by quantitative real-time PCR and found that only expression of ULBP2 is increased upon infection in both SVG-A and 293T cells (Fig 3C). Enhanced ULBP2 expression levels were also observed in JCPyV-infected human primary renal proximal tubule epithelial (HPRTE) cells, a primary target for JCPyV, upon secondary analysis of data generated in the laboratory of Walter J. Atwood [63] (Fig. 3D). Based on enhanced expression of ULBP2 on JCPyV-infected cells, we then performed co-culture assays in the presence of antibodies blocking the activating NKG2D receptor. Blockade of NKG2D resulted in decreased NK cell degranulation response to JCPyV-infected SVG-A cells (Fig 3E-F). Therefore, one of the mechanisms underlying NK cell-mediated responses to JCPyV-infected SVG-A cells relies on their recognition via NKG2D.

**Fig 3.**
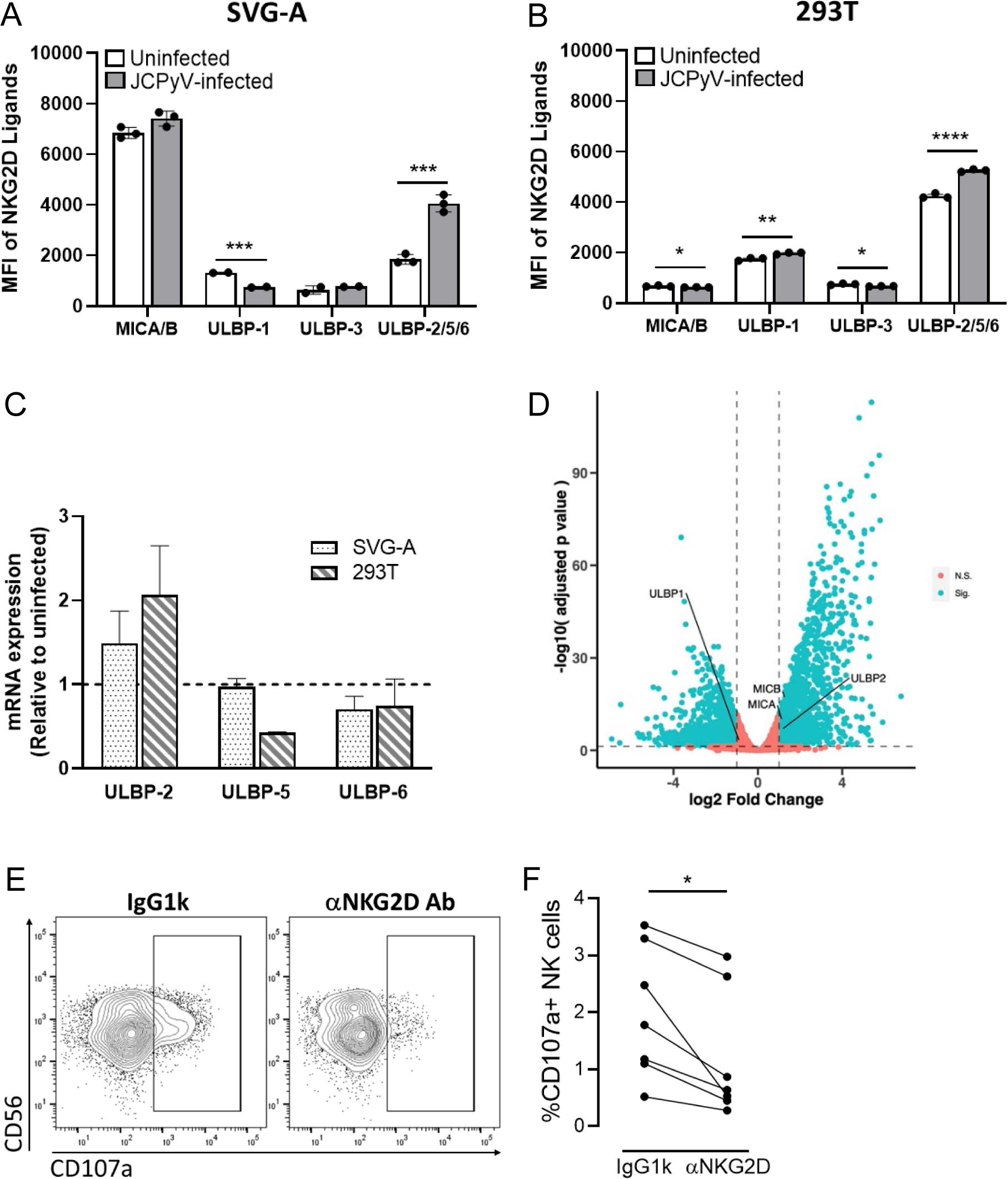
NKG2D and its ligand ULBP-2 play a key role in NK cell responses against JCPyV-infected cells. SVG-A (A) and 293T (B) cells were infected with M1-SVEΔ or cultured without virus and after 9 days, cell surface expression of ligands for NKG2D was assessed by flow cytometry. (C) RNA was extracted from uninfected and infected SVG-A and 293T cells to quantify mRNA expression levels using primers specific for each ULBP by TaqMan qPCR. (D) Volcano plot displaying fold-changes in transcripts levels for the indicated NKG2D ligands in HRPTE cells 9 days following infection with M1-SVEΔ compared to uninfected epithelial cells, using pre-computed log2 fold changes and p-values available here: https://www.ncbi.nlm.nih.gov/geo/query/acc.cgi?acc=GSE135833. N.S., not significant, defined as absolute value of log 2-fold change less than one and adjusted p-value less than 0.05; Sig., significant, defined as absolute value of log2 fold change greater than 1 and adjusted p-value less than 0.05. (E) Representative flow cytometry plots of CD107a+ NK cells following co-culture with SVG-A cells infected with SVE-Delta-M1 for 6h in the presence of isotype control (IgG1k) or blocking antibodies against NKG2D. (F) Compiled results for NK cells isolated from 7 healthy donors. *p<0.05; **p<0.01; *** p<0.001; **** p<0.0001.

## Discussion

Compelling evidence supports a crucial role for NK cells in the control of several major human viral infections as well as in shaping adaptive immune responses. However, there are many gaps in knowledge regarding the contribution of NK cells in the pathogenesis and progression of PML, the disease caused by JCPyV reactivation in the brain of immunocompromised individuals. Herein, we present the first evidence that JCPyV VP1-derived peptide stimulation and JCPyV infection both elicit robust NK cell function, suggesting that NK cells may significantly contribute to the cellular effector response to JCPyV. We also unveil mechanistic insights underlying immune recognition of a JCPyV-infected astrocyte cell line, which implicate NKG2D and its stress-induced ligand ULBP2.

While NKG2D engagement promotes potent and dominant NK cell activation, several viruses have evolved elaborate mechanisms to evade NKG2D-mediated recognition. For instance, studies have shown shedding of NKG2DL by HIV-infected CD4+ T cells, thereby promoting reduced expression of NKG2D on NK cells and impaired NKG2D-mediated NK cell responses [64, 65]. As PLWH represent the majority of PML patients, it is possible that impaired NKG2D signaling in PLWH is associated with poor NK cell-mediated control of JCPyV replication and development of PML. Future investigations are warranted to examine NK cell surface expression of NKG2D and plasma levels of soluble NKG2DL in PML patients, including both PLWH and HIV-negative patients.

Another reported immune escape mechanism exploited by viruses is the downregulation of NKG2DL expressed on infected cells. Interestingly, an elegant study previously reported that JCPyV encodes microRNAs that downregulate ULBP3, thereby reducing NK cell-mediated recognition of polyomaviruses-infected cells [32]. In our system, we did not observe significant changes in ULBP3 expression upon infection. This discrepancy may be explained by the different cell lines and JCPyV strain used, as well as the later time point for the assessment of NKG2DL expression in our experiments compared to the studies conducted by Bauman et al. Many immortalized cell lines trigger potent NK cell cytotoxicity because they express ligands for activating NK cell receptors or lack HLA-Class I ligands for inhibitory NK cell receptors [66]. SVG-A cells were selected for our experiments because infection of these cells is a long standing and well-established *in vitro* system to study JCPyV [103, 104] and compared to other cell lines permissive for JCPyV infection such as 293T, 293TT and C33A cells, uninfected SVG-A cells elicited only background levels of NK cell responses (data not shown). Nevertheless, altogether, these findings are consistent with differential expression of the various NKG2DL in different primary cells or cell lines and all support a significant role for the NKG2D pathway in the recognition of JCPyV-infected cells by NK cells. This data opens avenues for therapeutics against PML based on existing strategies currently in development targeting NKG2D [45–48].

Finally, NK cell responses to JCPyV most likely do not solely depend on the NKG2D/NKG2DL axis, and other pathways may be additive or alternative to this mechanism. Several immunotherapeutic strategies to treat PML have been evaluated in individual case reports and small patient series, including stimulation of lymphocytes using IL-7 [29, 67–72] and IL-15 [30]. These approaches have been associated with improved PML outcome in subsets of patients and since IL-7 and IL-15 are known to activate NK cells, their contribution to the clinical outcome of these therapies cannot be excluded. Notably, successful treatment with IL-15 superagonist in a PML patient with allogeneic stem cell transplant was associated with a rise in peripheral blood natural killer (NK) cells but not in CD3+ T cells [30]. Overall, rationally optimizing therapies under investigation by targeting bulk or subsets of NK cells may offer novel immunotherapeutic approaches against PML.

Our data also show NK cell activation, and particularly IFN-y production, upon stimulation with VP1-derived peptides. Interestingly, there was no positive correlation between the proportions of IL-2+ T cells and those of IFN-y+ NK cells (data not shown), indicating that in this assay, NK cell responses may not mainly rely on IL-2 produced by JCPyV-specific memory T cells. NK cell activation could be directly triggered by VP1 peptides presented by HLA class I that either disrupt the interaction between HLA class I molecules and their inhibitory ligand on NK cells, or engage NK cell activating receptors. Further investigations to explore the role played by inhibitory and activating killer cell immunoglobulin like receptors (KIRs) or by CD94-NKG2A/C heterodimers in these responses are warranted.

Collectively, these findings suggest that the loss of cellular immunity associated with enhanced JCPyV replication and progression towards PML may encompass impaired NK cell function, and that boosting NK cell activity may potentiate the overall immune control of JCPyV. To our knowledge, these results provide the first evidence for a direct effect mediated by NK cells against JCPyV and have important implications for the design of future immunotherapeutic interventions aimed at enhancing JCPyV immunity in immunocompromised individuals.

## Materials and Methods

### Human Subjects

Deidentified and coded blood samples from healthy donors used in this study were collected under IRB-approved protocols and delivered to us by Research Blood Components, LLC (Watertown, Massachusetts). Frozen PBMC vials from healthy donors were purchased from STEMCELL Technologies. Beth Israel Deaconess Medical Center institutional review board approved this study, and all subjects gave written informed consent.

### Analysis of primary NK cell responses to JCPyV VP1-derived peptides by intracellular cytokine staining

To measure NK and T cell responses to the JCPyV capsid VP1, cryopreserved PBMCs from healthy donors that tested seropositive for JCPyV were thawed and directly stimulated with 2μg/mL of sequential peptide pools consisting of 15-mer sequences with 11 aa overlap covering the whole JCPyV VP1 sequence and combined into 2 pools (VP1 1-93 + VP1 97-157 (n=48); VP1 161-253 + VP1 257-341 (n=49). PBMC were co-cultured with peptides at 37°C for 6h with CD107a BV786 (BD Biosciences, H4A3) and 1μg/mL of CD28/CD49d costimulatory reagent (BD Biosciences). 1μL/mL GolgiPlug (BD Biosciences) and 0.7μL/mL GolgiStop (BD Biosciences) were added for the last 2h of incubation. At the end of the incubation, cells were stained first with the LIVE/DEAD® Fixable Blue Dead Cell Stain Kit (Invitrogen), then with BD Biosciences CD3 BV510 (UCHT1), CD14 BV421 (M5E2), CD19 BV421 (HIB19), CD16 APC-Cy7 (3G8), CD56 BV605 (NCAM16.2), CD4 BUV395 (L200) and CD8 APC (SK1), and finally fixed, permeabilized (Thermofisher Fix and Perm) and stained with BD Biosciences IFN-γ FITC (B27) and IL-2 PE-CF594 (5344.11) antibodies to detect intracellular cytokines. In all assays described above, incubation in the presence of 5 μg/mL of phytohemagglutinin (PHA) was used as positive control and unstimulated cells served as negative controls and for background subtraction. A Fluorescence Minus One (FMO) control and PHA-stimulated PBMCs were used to set the gates for positive cytokine responses. Acquisition of data was performed on a BD LSRII instrument (BD Biosciences). Data was analyzed using Flow Jo v.10.8.1.

### JCPyV infection of SVG-A cells

SVG-A cells were grown in EMEM supplemented with 2% FBS and 2% Penicillin/streptomycin. 2-3 days prior infection, SVG-A cells were plated into two T-75 flasks with 0.7M cells in each flask. After 2-3 days, one flask was infected with JCPyV M1-SVEΔ (5.9e9 viral genome/mL) at a 1:100 dilution and one flask left uninfected. Cells in both flasks were cultured for a total of 9-11 days at 37ºC, with addition of fresh media one-week post-infection. Percentages of infected SVG-A cells were determined prior to functional assays by intracellular staining using Thermofisher antibodies against JCPyV VP1 DyLight488 (PAB597) and Large T antigen DyLight594 (PAB2003) on the BDLSRFortessa X14.

### Killing assay

NK Cells were enriched from freshly isolated PBMC using the EasySep™ Human NK Cell Isolation Kit (STEMCELL Technologies). NK cells were then counted and resuspended at 1M/mL in RPMI-1640 supplemented with 10% fetal bovine serum, 2 mM L-glutamine, 100 μg/mL streptomycin and 100 U/mL penicillin. Based on the number of infected SVG-A determined beforehand on the same day by flow cytometry, SVG-A cells and NK cells were combined at an E:T ratio of 10:1 (NK cells: infected SVG-A cells). To complement measures of direct NK cell-mediated killing, CD107a BV786 was also added at a 5μL/mL concentration and served as a surrogate marker of degranulation. In a subset of experiments where only CD107a upregulation but not killing was measured, 1μL/mL GolgiPlug (BD Biosciences) and 0.7μL/mL GolgiStop (BD Biosciences) were added for the whole incubation (S2 Fig). For blockade experiments, purified NK cells were incubated for 10 minutes at room temperature with 2.5µg of Human BD Fc Block followed by addition of 10μg/mL of anti-NKG2D purified antibody (BD Biosciences, clone 1D11) or isotype control antibodies prior to addition of SVG-A cells. Co-cultures were incubated at 37ºC for 6hr. At the end of the incubation, cells were stained first with the LIVE/DEAD® Fixable Aqua Dead Cell Stain Kit (Invitrogen), then with BD Biosciences CD3 A700 (UCHT1), CD16 APC-Cy7 (3G8), CD56 BV605 (NCAM16.2), NKG2D APC (1D11) and Beckman Coulter NKG2A PE-Cy7 (Z199), fixed, permeabilized (Thermofisher Fix and Perm), and stained with Thermofisher VP1 DyLight488 (PAB597) and Large T antigen DyLight594 (PAB2003) to quantify SVG-A infected cells. Acquisition of data was performed on a BD FACSymphony A5. Data was analyzed using Flow Jo v.10.8.1.

### Anti-JCPyV IgG ELISA

JCPyV capsid protein VP1 was obtained from Abcam (AB74569). Endpoint titer dilution ELISA was performed to determine the JCPyV serostatus of all healthy donors included in our study as previously published [73]. Briefly, polystyrene ELISA plates were coated overnight with 1ug/well of JCPyV VP1 protein. After blocking with 1% casein (Thermofisher Blocker™ Casein) and washing with PBS containing 0.05% Tween 20, plasmas were added in serial dilutions and incubated for 1h at room temperature. The plates were then washed three times with PBS containing 0.05% Tween 20 and incubated for 1 hr with a dilution of a 1/1,000 horseradish peroxidase (HRP)-conjugated goat anti-human secondary antibody (Jackson ImmunoResearch Laboratories), developed with TMB Peroxidase Substrate (SeraCare), and stopped by addition of stopping solution (SeraCare), and analyzed at 450/550nm with Spectramax Plus ELISA plate reader using Softmas Pro 4.7.1 software. ELISA endpoint titers were defined as the highest reciprocal plasma dilution that yielded an absorbance > 2-fold over background values.

### Quantification of NKG2DL by flow cytometry

SVG-A and 293T cells were infected with JCPyV M1-SVEΔ for 9-11 days and stained first with the LIVE/DEAD® Fixable Aqua Dead Cell Stain Kit (Invitrogen), then cells were split and stained in parallel with one of the following antibodies: R&D Systems ULBP1 PE (170818), ULBP3 PE (166510), ULBP-2/5/6 PE (165903) or BD Biosciences MICA/B PE (6D4) prior to be fixed, permeabilized and stained with Thermofisher VP1 DyLight488 (PAB597) and Large T antigen DyLight594 (PAB2003)with. Uninfected SVG-A and 293T cells as well as Raji and K562 cells known to express low and high levels of NKG2DL, respectively [66], were included as controls.

### Quantification of NKG2DL by qPCR

Two and a half μg of total RNA isolated using Qiagen RNeasy kit, was used for cDNA first-strand synthesis in a 20-μL reaction volume using Thermofisher SuperScript™ VILO™ cDNA Synthesis Kit. Real-time quantitative PCR was performed using the QuantStudio™ 3 Flex Real-Time PCR System (Applied Biosystems) and TaqMan™ Fast Advanced Master Mix. cDNA was amplified with specific commercially available primers for ULBP2 (Hs00607609_mH), ULBP3 (Hs00225909_m1), ULBP5 (Hs01584111_mH), ULBP6 (Hs04194671_s1) and beta actin (Hs99999903_m1) with TaqMan MGB probes all conjugated with fluorochrome FAM (Thermofisher). The cycling conditions were 50°C for 2 minutes, polymerase activation at 95°C for 20 seconds, followed by 40 cycles of 95°C for 1 second, and 60°C for 20 seconds. Data were analyzed using Design & Analysis Software (version 2.6.0, Applied Biosystems) and R version 4.2.3 (https://www.Rproject.org).

### Statistical Analyses

All bars in scatter dot plots represent median values. Column bar graphs represent mean ± SEM. Statistical analyses were performed using the GraphPad Prism software version 9.5.0 The non-parametric Wilcoxon signed-rank test was used to assess differences in cytokine production or CD107a upregulation in response to VP1-derived peptides, to JCPyV infection and in the presence or absence of blocking antibodies, or differences in proportions of JCPyV-infected cells in the presence or absence of NK cells. Unpaired *t* tests were used to compare expression of NKG2DL between uninfected and JCPyV-infected cells. P-values of <0.05 were considered significant.

### RNA transcriptome analysis

Transcriptome profiling of infected HRPTE cells and data analysis were performed as previously described [63]. Deposited data including pre-computed log2 fold changes and p-values were used to generate volcano plots using ggplot2 v. 3.3.2. and are available here: https://www.ncbi.nlm.nih.gov/geo/query/acc.cgi?acc=GSE135833.

## Acknowledgments

We would like to thank R. Keith Reeves for his valuable feedback and helpful comments on this manuscript.

## Supporting information captions

**S1 Fig.**
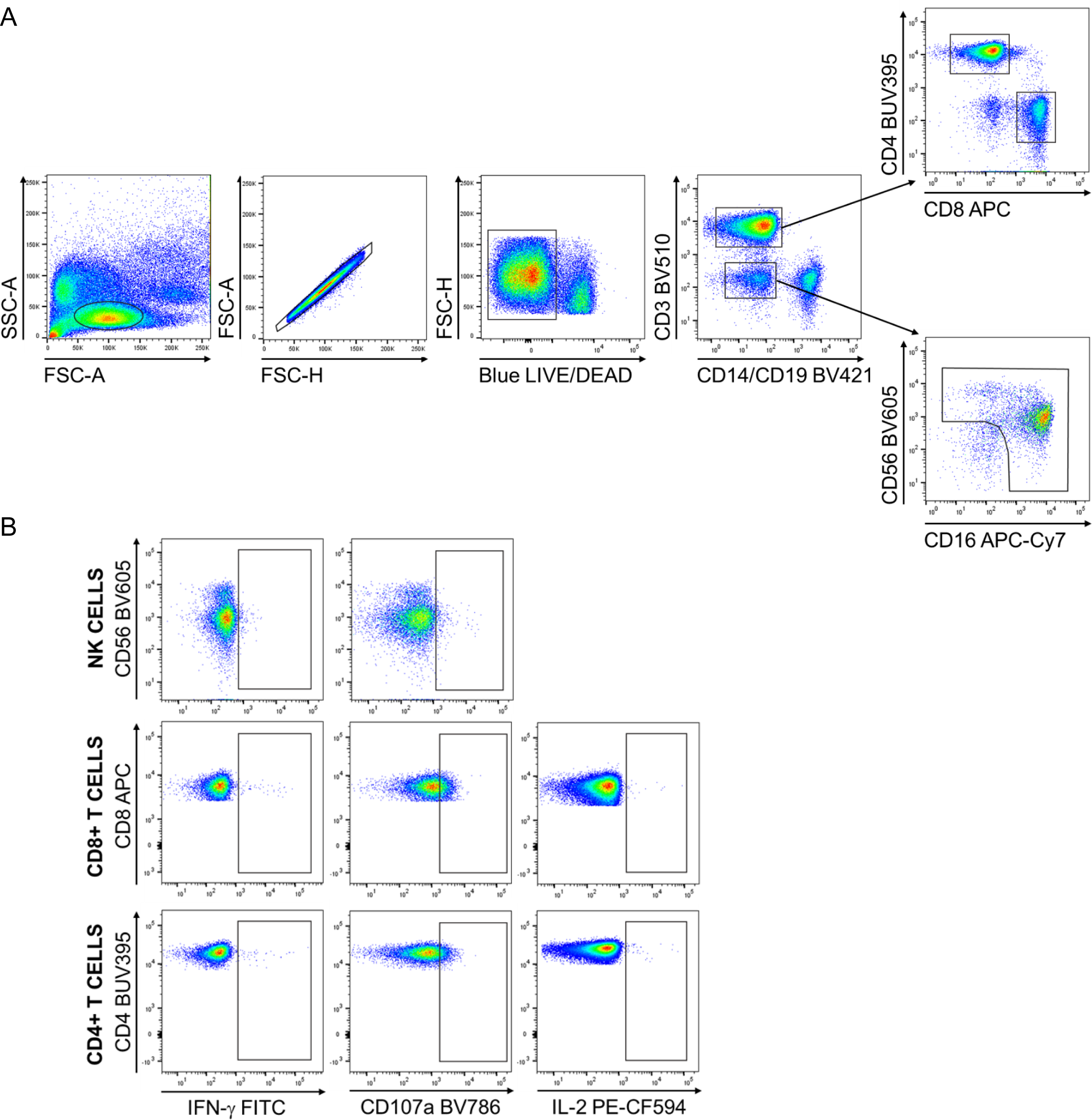
Gating strategy to measure NK and T cell responses to JCPyV VP1 by ICS. (Related to Fig 1). (A) Flow cytometry gating strategy to analyze intracellular expression of CD107a and IFN-y on NK, CD4+ T and CD8+ T cells as well as IL-2 on CD4+ T and CD8+ T cells. Gates are set to exclude doublets, dead cells, CD14+ and CD19+ cells. (B) Representative primary flow cytometry plots showing CD107a upregulation as well as IFN-y and IL-2 production by NK cells, CD8+T and CD4+ T cells as indicated following stimulation with the JCPyV peptide pool covering VP1 161-253 + VP1 257-341. Expression of functional markers on various lymphocyte subsets was defined by using appropriate unstimulated controls.

**S2 Fig.**
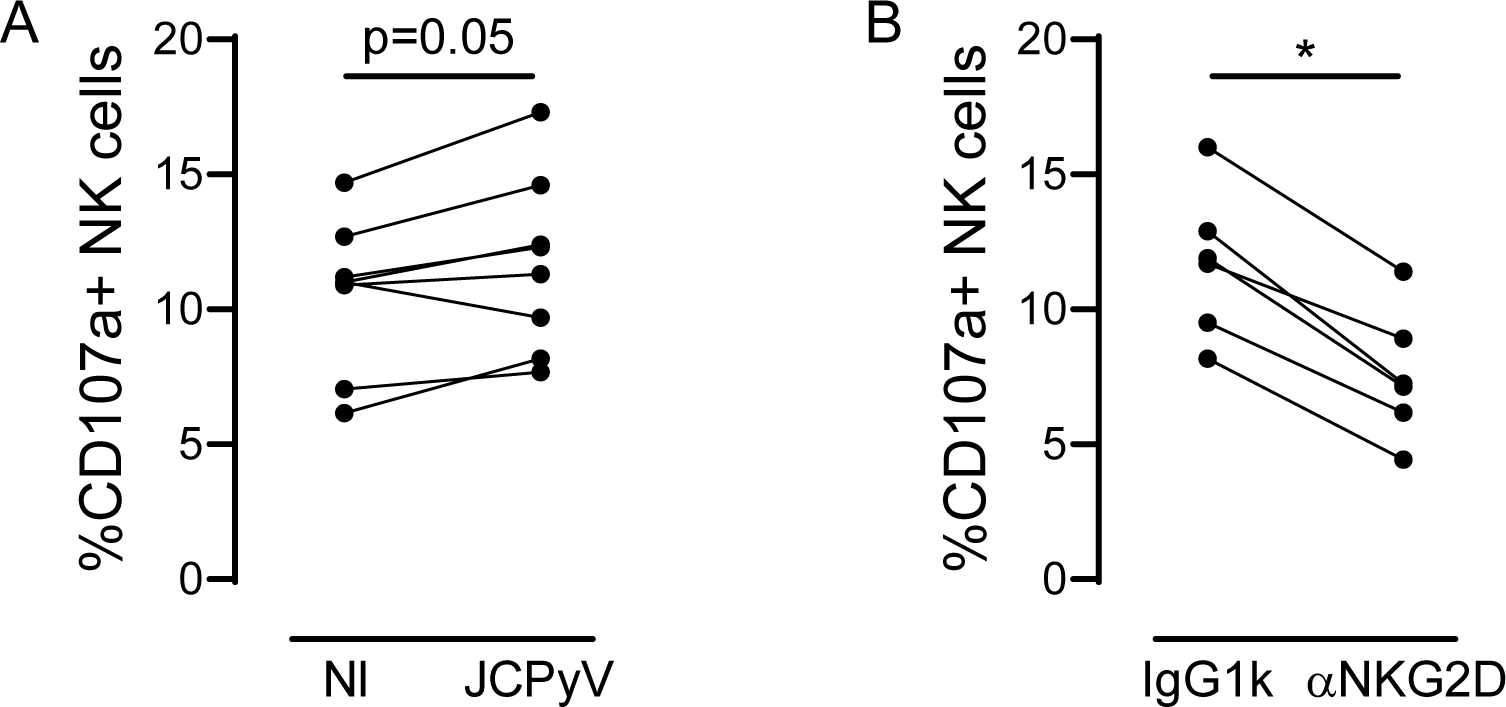
NKG2D blockade results in decreased NK cell response against JCPyV-infected cells. (Related to Fig 2D and 3E). (A) Proportions of CD107a+ NK cells from 8 healthy donors co-cultured with uninfected (NI) SVG-A cells or SVG-A cells infected with JCPyV M1-SVEΔ. Proportions of CD107a+ NK cells from 6 healthy donors following co-culture with SVG-A cells infected with M1-SVEΔ in the presence of isotype control (IgG1k) or blocking antibodies against NKG2D. NK cells and SVG-A cells were co-cultured for 6h in the presence of conjugated anti-CD107a antibodies and 1μL/mL GolgiPlug (BD Biosciences) and 0.7μL/mL GolgiStop (BD Biosciences).**p<0.01; **** p<0.0001.

## References

1. Knowles WA, Pipkin P, Andrews N, Vyse A, Minor P, Brown DW, et al. Population-based study of antibody to the human polyomaviruses BKV and JCV and the simian polyomavirus SV40. J Med Virol. 2003;71(1):115–23. Epub 2003/07/15. doi: 10.1002/jmv.10450. PubMed PMID: 12858417.

2. Kitamura T, Sugimoto C, Kato A, Ebihara H, Suzuki M, Taguchi F, et al. Persistent JC virus (JCV) infection is demonstrated by continuous shedding of the same JCV strains. J Clin Microbiol. 1997;35(5):1255–7. Epub 1997/05/01. doi: 10.1128/jcm.35.5.1255-1257.1997. PubMed PMID: 9114418; PubMed Central PMCID: PMCPMC232740.

3. Tan CS, Dezube BJ, Bhargava P, Autissier P, Wuthrich C, Miller J, et al. Detection of JC virus DNA and proteins in the bone marrow of HIV-positive and HIV-negative patients: implications for viral latency and neurotropic transformation. J Infect Dis. 2009;199(6):881–8. Epub 2009/05/13. doi: 10.1086/597117. PubMed PMID: 19434914; PubMed Central PMCID: PMCPMC2893283.

4. Monaco MC, Atwood WJ, Gravell M, Tornatore CS, Major EO. JC virus infection of hematopoietic progenitor cells, primary B lymphocytes, and tonsillar stromal cells: implications for viral latency. J Virol. 1996;70(10):7004–12. Epub 1996/10/01. doi: 10.1128/JVI.70.10.7004-7012.1996. PubMed PMID: 8794345; PubMed Central PMCID: PMCPMC190751.

5. Cortese I, Reich DS, Nath A. Progressive multifocal leukoencephalopathy and the spectrum of JC virus-related disease. Nat Rev Neurol. 2021;17(1):37–51. Epub 2020/11/22. doi: 10.1038/s41582-020-00427-y. PubMed PMID: 33219338; PubMed Central PMCID: PMCPMC7678594.

6. Tan CS, Koralnik IJ. Progressive multifocal leukoencephalopathy and other disorders caused by JC virus: clinical features and pathogenesis. Lancet Neurol. 2010;9(4):425–37. Epub 2010/03/20. doi: 10.1016/S1474-4422(10)70040-5. PubMed PMID: 20298966; PubMed Central PMCID: PMC2880524.

7. Caniglia EC, Cain LE, Justice A, Tate J, Logan R, Sabin C, et al. Antiretroviral penetration into the CNS and incidence of AIDS-defining neurologic conditions. Neurology. 2014;83(2):134–41. Epub 2014/06/08. doi: 10.1212/WNL.0000000000000564. PubMed PMID: 24907236; PubMed Central PMCID: PMC4117168.

8. Clifford DB, Yiannoutsos C, Glicksman M, Simpson DM, Singer EJ, Piliero PJ, et al. HAART improves prognosis in HIV-associated progressive multifocal leukoencephalopathy. Neurology. 1999;52(3):623–5. Epub 1999/02/20. PubMed PMID: 10025799.

9. Lane MA, Renga V, Pachner AR, Cohen JA. Late Occurrence of PML in a Patient Treated for Lymphoma with Immunomodulatory Chemotherapies, Bendamustine, Rituximab, and Ibritumomab Tiuxetan. Case Rep Neurol Med. 2015;2015:892047. Epub 2015/02/24. doi: 10.1155/2015/892047. PubMed PMID: 25705531; PubMed Central PMCID: PMC4326342.

10. Isidoro L, Pires P, Rito L, Cordeiro G. Progressive multifocal leukoencephalopathy in a patient with chronic lymphocytic leukaemia treated with alemtuzumab. BMJ Case Rep. 2014;2014. Epub 2014/01/10. doi: 10.1136/bcr-2013-201781. PubMed PMID: 24403383.

11. Boster AL, Nicholas JA, Topalli I, Kisanuki YY, Pei W, Morgan-Followell B, et al. Lessons learned from fatal progressive multifocal leukoencephalopathy in a patient with multiple sclerosis treated with natalizumab. JAMA Neurol. 2013;70(3):398–402. Epub 2013/01/23. doi: 10.1001/jamaneurol.2013.1960. PubMed PMID: 23338729.

12. Bloomgren G, Richman S, Hotermans C, Subramanyam M, Goelz S, Natarajan A, et al. Risk of natalizumab-associated progressive multifocal leukoencephalopathy. N Engl J Med. 2012;366(20):1870–80. Epub 2012/05/18. doi: 10.1056/NEJMoa1107829. PubMed PMID: 22591293.

13. Clifford DB, Nath A, Cinque P, Brew BJ, Zivadinov R, Gorelik L, et al. A study of mefloquine treatment for progressive multifocal leukoencephalopathy: results and exploration of predictors of PML outcomes. Journal of neurovirology. 2013. Epub 2013/06/05. doi: 10.1007/s13365-013-0173-y. PubMed PMID: 23733308.

14. Cettomai D, McArthur JC. Mirtazapine use in human immunodeficiency virus-infected patients with progressive multifocal leukoencephalopathy. Arch Neurol. 2009;66(2):255–8. doi: 10.1001/archneurol.2008.557. PubMed PMID: 19204164.

15. Hall CD, Dafni U, Simpson D, Clifford D, Wetherill PE, Cohen B, et al. Failure of cytarabine in progressive multifocal leukoencephalopathy associated with human immunodeficiency virus infection. AIDS Clinical Trials Group 243 Team. N Engl J Med. 1998;338(19):1345-51. doi: 10.1056/NEJM199805073381903. PubMed PMID: 9571254.

16. Marra CM, Rajicic N, Barker DE, Cohen BA, Clifford D, Donovan Post MJ, et al. A pilot study of cidofovir for progressive multifocal leukoencephalopathy in AIDS. Aids. 2002;16(13):1791–7.

17. Tan K, Roda R, Ostrow L, McArthur J, Nath A. PML-IRIS in patients with HIV infection: clinical manifestations and treatment with steroids. Neurology. 2009;72(17):1458–64. Epub 2009/01/09. doi: 10.1212/01.wnl.0000343510.08643.74. PubMed PMID: 19129505; PubMed Central PMCID: PMCPMC2677476.

18. Tan IL, McArthur JC, Clifford DB, Major EO, Nath A. Immune reconstitution inflammatory syndrome in natalizumab-associated PML. Neurology. 2011;77(11):1061–7. Epub 2011/08/13. doi: 10.1212/WNL.0b013e31822e55e7. PubMed PMID: 21832229; PubMed Central PMCID: PMCPMC3174071.

19. N’Gbo N’gbo Ikazabo R, Mostosi C, Quivron B, Delberghe X, El Hafsi K, Lysandropoulos AP. Immune-reconstitution Inflammatory Syndrome in Multiple Sclerosis Patients Treated With Natalizumab: A Series of 4 Cases. Clin Ther. 2016;38(3):670–5. Epub 2016/02/10. doi: 10.1016/j.clinthera.2016.01.010. PubMed PMID: 26856928.

20. Vendrely A, Bienvenu B, Gasnault J, Thiebault JB, Salmon D, Gray F. Fulminant inflammatory leukoencephalopathy associated with HAART-induced immune restoration in AIDS-related progressive multifocal leukoencephalopathy. Acta Neuropathol. 2005;109(4):449–55. Epub 2005/03/02. doi: 10.1007/s00401-005-0983-y. PubMed PMID: 15739098.

21. Gheuens S, Bord E, Kesari S, Simpson DM, Gandhi RT, Clifford DB, et al. Role of CD4+ and CD8+ T-cell responses against JC virus in the outcome of patients with progressive multifocal leukoencephalopathy (PML) and PML with immune reconstitution inflammatory syndrome. J Virol. 2011;85(14):7256–63. Epub 2011/05/06. doi: 10.1128/JVI.02506-10. PubMed PMID: 21543472; PubMed Central PMCID: PMC3126613.

22. Du Pasquier RA, Kuroda MJ, Zheng Y, Jean-Jacques J, Letvin NL, Koralnik IJ. A prospective study demonstrates an association between JC virus-specific cytotoxic T lymphocytes and the early control of progressive multifocal leukoencephalopathy. Brain. 2004;127(Pt 9):1970–8. Epub 2004/06/25. doi: 10.1093/brain/awh215. PubMed PMID: 15215217.

23. Du Pasquier RA, Schmitz JE, Jean-Jacques J, Zheng Y, Gordon J, Khalili K, et al. Detection of JC virus-specific cytotoxic T lymphocytes in healthy individuals. J Virol. 2004;78(18):10206–10. Epub 2004/08/28. doi: 10.1128/JVI.78.18.10206-10210.2004. PubMed PMID: 15331755; PubMed Central PMCID: PMCPMC514969.

24. Du Pasquier RA, Stein MC, Lima MA, Dang X, Jean-Jacques J, Zheng Y, et al. JC virus induces a vigorous CD8+ cytotoxic T cell response in multiple sclerosis patients. J Neuroimmunol. 2006;176(1-2):181–6. Epub 2006/06/06. doi: 10.1016/j.jneuroim.2006.04.003. PubMed PMID: 16750575.

25. Koralnik IJ, Du Pasquier RA, Letvin NL. JC virus-specific cytotoxic T lymphocytes in individuals with progressive multifocal leukoencephalopathy. J Virol. 2001;75(7):3483–7. Epub 2001/03/10. doi: 10.1128/JVI.75.7.3483-3487.2001. PubMed PMID: 11238876; PubMed Central PMCID: PMCPMC114143.

26. Ferenczy MW, Marshall LJ, Nelson CD, Atwood WJ, Nath A, Khalili K, et al. Molecular biology, epidemiology, and pathogenesis of progressive multifocal leukoencephalopathy, the JC virus-induced demyelinating disease of the human brain. Clin Microbiol Rev. 2012;25(3):471–506. Epub 2012/07/06. doi: 10.1128/CMR.05031-11. PubMed PMID: 22763635; PubMed Central PMCID: PMCPMC3416490.

27. Khanna N, Wolbers M, Mueller NJ, Garzoni C, Du Pasquier RA, Fux CA, et al. JC virus-specific immune responses in human immunodeficiency virus type 1 patients with progressive multifocal leukoencephalopathy. J Virol. 2009;83(9):4404–11. Epub 2009/02/13. doi: 10.1128/JVI.02657-08. PubMed PMID: 19211737; PubMed Central PMCID: PMCPMC2668450.

28. Muftuoglu M, Olson A, Marin D, Ahmed S, Mulanovich V, Tummala S, et al. Allogeneic BK Virus-Specific T Cells for Progressive Multifocal Leukoencephalopathy. N Engl J Med. 2018;379(15):1443–51. Epub 2018/10/12. doi: 10.1056/NEJMoa1801540. PubMed PMID: 30304652; PubMed Central PMCID: PMCPMC6283403.

29. Alstadhaug KB, Croughs T, Henriksen S, Leboeuf C, Sereti I, Hirsch HH, et al. Treatment of progressive multifocal leukoencephalopathy with interleukin 7. JAMA Neurol. 2014;71(8):1030–5. Epub 2014/07/01. doi: 10.1001/jamaneurol.2014.825. PubMed PMID: 24979548.

30. Oza A, Rettig MP, Powell P, O’Brien K, Clifford DB, Ritchey J, et al. Interleukin-15 superagonist (N-803) treatment of PML and JCV in a post-allogeneic hematopoietic stem cell transplant patient. Blood Adv. 2020;4(11):2387–91. Epub 2020/06/03. doi: 10.1182/bloodadvances.2019000664. PubMed PMID: 32484854; PubMed Central PMCID: PMCPMC7284083.

31. Gheuens S, Bord E, Kesari S, Simpson DM, Gandhi RT, Clifford DB, et al. Role of CD4+ and CD8+ T-cell responses against JC virus in the outcome of patients with progressive multifocal leukoencephalopathy (PML) and PML with immune reconstitution inflammatory syndrome. J Virol. 2011;85(14):7256–63. Epub 2011/05/06. doi: 10.1128/JVI.02506-10JVI.02506-10 [pii]. PubMed PMID: 21543472; PubMed Central PMCID: PMC3126613.

32. Ray U, Cinque P, Gerevini S, Longo V, Lazzarin A, Schippling S, et al. JC polyomavirus mutants escape antibody-mediated neutralization. Science translational medicine. 2015;7(306):306ra151. Epub 2015/09/25. doi: 10.1126/scitranslmed.aab1720. PubMed PMID: 26400912.

33. Tan CS, Ghofrani J, Geiger E, Koralnik IJ, Jost S. Brief Report: Decreased JC Virus-Specific Antibody-Dependent Cellular Cytotoxicity in HIV-Seropositive PML Survivors. J Acquir Immune Defic Syndr. 2019;82(2):220–4. Epub 2019/09/13. doi: 10.1097/QAI.000000000000210500126334-201910010-00017 [pii]. PubMed PMID: 31513076; PubMed Central PMCID: PMC7179760.

34. Bauman Y, Nachmani D, Vitenshtein A, Tsukerman P, Drayman N, Stern-Ginossar N, et al. An identical miRNA of the human JC and BK polyoma viruses targets the stress-induced ligand ULBP3 to escape immune elimination. Cell host & microbe. 2011;9(2):93–102. Epub 2011/02/16. doi: 10.1016/j.chom.2011.01.008. PubMed PMID: 21320692.

35. Lanier LL. NK cell recognition. Annu Rev Immunol. 2005;23:225-74. PubMed PMID: 15771571.

36. Diefenbach A, Jensen ER, Jamieson AM, Raulet DH. Rae1 and H60 ligands of the NKG2D receptor stimulate tumour immunity. Nature. 2001;413(6852):165-71. Epub 2001/09/15. doi: 10.1038/35093109. PubMed PMID: 11557981; PubMed Central PMCID: PMCPMC3900321.

37. Groh V, Rhinehart R, Randolph-Habecker J, Topp MS, Riddell SR, Spies T. Costimulation of CD8alphabeta T cells by NKG2D via engagement by MIC induced on virus-infected cells. Nat Immunol. 2001;2(3):255–60. Epub 2001/02/27. doi: 10.1038/85321. PubMed PMID: 11224526.

38. Groh V, Bahram S, Bauer S, Herman A, Beauchamp M, Spies T. Cell stress-regulated human major histocompatibility complex class I gene expressed in gastrointestinal epithelium. Proc Natl Acad Sci U S A. 1996;93(22):12445–50. Epub 1996/10/29. doi: 10.1073/pnas.93.22.12445. PubMed PMID: 8901601; PubMed Central PMCID: PMCPMC38011.

39. Cao W, Xi X, Wang Z, Dong L, Hao Z, Cui L, et al. Four novel ULBP splice variants are ligands for human NKG2D. Int Immunol. 2008;20(8):981–91. Epub 2008/06/12. doi: 10.1093/intimm/dxn057. PubMed PMID: 18544572.

40. Bauer S, Groh V, Wu J, Steinle A, Phillips JH, Lanier LL, et al. Activation of NK cells and T cells by NKG2D, a receptor for stress-inducible MICA. Science. 1999;285(5428):727-9. Epub 1999/07/31. doi: 10.1126/science.285.5428.727. PubMed PMID: 10426993.

41. Lanier LL. NKG2D Receptor and Its Ligands in Host Defense. Cancer Immunol Res. 2015;3(6):575–82. Epub 2015/06/05. doi: 10.1158/2326-6066.CIR-15-0098. PubMed PMID: 26041808; PubMed Central PMCID: PMCPMC4457299.

42. Fang M, Lanier LL, Sigal LJ. A role for NKG2D in NK cell-mediated resistance to poxvirus disease. PLoS Pathog. 2008;4(2):e30. Epub 2008/02/13. doi: 10.1371/journal.ppat.0040030. PubMed PMID: 18266471; PubMed Central PMCID: PMCPMC2233669 have licensed intellectual property rights relating to NKG2D for commercial applications. The authors declare that no other competing interests exist.

43. Etzioni A, Eidenschenk C, Katz R, Beck R, Casanova JL, Pollack S. Fatal varicella associated with selective natural killer cell deficiency. J Pediatr. 2005;146(3):423–5. Epub 2005/03/10. doi: 10.1016/j.jpeds.2004.11.022. PubMed PMID: 15756234.

44. Biron CA, Byron KS, Sullivan JL. Severe herpesvirus infections in an adolescent without natural killer cells. N Engl J Med. 1989;320(26):1731–5. PubMed PMID: 2543925.

45. Zhang C, Roder J, Scherer A, Bodden M, Pfeifer Serrahima J, Bhatti A, et al. Bispecific antibody-mediated redirection of NKG2D-CAR natural killer cells facilitates dual targeting and enhances antitumor activity. J Immunother Cancer. 2021;9(10). Epub 2021/10/03. doi: 10.1136/jitc-2021-002980. PubMed PMID: 34599028; PubMed Central PMCID: PMCPMC8488744.

46. Baumeister SH, Murad J, Werner L, Daley H, Trebeden-Negre H, Gicobi JK, et al. Phase I Trial of Autologous CAR T Cells Targeting NKG2D Ligands in Patients with AML/MDS and Multiple Myeloma. Cancer Immunol Res. 2019;7(1):100–12. Epub 2018/11/07. doi: 10.1158/2326-6066.CIR-18-0307. PubMed PMID: 30396908; PubMed Central PMCID: PMCPMC7814996.

47. Fuertes MB, Domaica CI, Zwirner NW. Leveraging NKG2D Ligands in Immuno-Oncology. Front Immunol. 2021;12:713158. Epub 2021/08/17. doi: 10.3389/fimmu.2021.713158. PubMed PMID: 34394116; PubMed Central PMCID: PMCPMC8358801.

48. Curio S, Jonsson G, Marinovic S. A summary of current NKG2D-based CAR clinical trials. Immunother Adv. 2021;1(1):ltab018. Epub 2021/10/05. doi: 10.1093/immadv/ltab018. PubMed PMID: 34604863; PubMed Central PMCID: PMCPMC8480431.

49. de Haas M, Koene HR, Kleijer M, de Vries E, Simsek S, van Tol MJ, et al. A triallelic Fc gamma receptor type IIIA polymorphism influences the binding of human IgG by NK cell Fc gamma RIIIa. J Immunol. 1996;156(8):2948–55. Epub 1996/04/15. PubMed PMID: 8609432.

50. Jawahar S, Moody C, Chan M, Finberg R, Geha R, Chatila T. Natural Killer (NK) cell deficiency associated with an epitope-deficient Fc receptor type IIIA (CD16-II). Clin Exp Immunol. 1996;103(3):408–13. Epub 1996/03/01. doi: 10.1111/j.1365-2249.1996.tb08295.x. PubMed PMID: 8608639; PubMed Central PMCID: PMC2200361.

51. de Vries E, Koene HR, Vossen JM, Gratama JW, von dem Borne AE, Waaijer JL, et al. Identification of an unusual Fc gamma receptor IIIa (CD16) on natural killer cells in a patient with recurrent infections. Blood. 1996;88(8):3022–7. Epub 1996/10/15. PubMed PMID: 8874200.

52. Yoon JC, Yang CM, Song Y, Lee JM. Natural killer cells in hepatitis C: Current progress. World J Gastroenterol. 2016;22(4):1449–60. Epub 2016/01/29. doi: 10.3748/wjg.v22.i4.1449. PubMed PMID: 26819513; PubMed Central PMCID: PMC4721979.

53. Florez-Alvarez L, Hernandez JC, Zapata W. NK Cells in HIV-1 Infection: From Basic Science to Vaccine Strategies. Front Immunol. 2018;9:2290. Epub 2018/11/06. doi: 10.3389/fimmu.2018.02290. PubMed PMID: 30386329; PubMed Central PMCID: PMC6199347.

54. Schultz-Cherry S. Role of NK cells in influenza infection. Curr Top Microbiol Immunol. 2015;386:109–20. Epub 2014/07/06. doi: 10.1007/82_2014_403. PubMed PMID: 24992894.

55. Fadda L, Borhis G, Ahmed P, Cheent K, Pageon SV, Cazaly A, et al. Peptide antagonism as a mechanism for NK cell activation. Proceedings of the National Academy of Sciences of the United States of America. 2010;107(22):10160–5. Epub 2010/05/05. doi: 10.1073/pnas.0913745107. PubMed PMID: 20439706; PubMed Central PMCID: PMC2890497.

56. Holzemer A, Garcia-Beltran WF, Altfeld M. Natural Killer Cell Interactions with Classical and Non-Classical Human Leukocyte Antigen Class I in HIV-1 Infection. Front Immunol. 2017;8:1496. Epub 2017/12/01. doi: 10.3389/fimmu.2017.01496. PubMed PMID: 29184550; PubMed Central PMCID: PMCPMC5694438.

57. Jost S, Tomezsko PJ, Rands K, Toth I, Lichterfeld M, Gandhi RT, et al. CD4+ T-cell help enhances NK cell function following therapeutic HIV-1 vaccination. Journal of virology. 2014;88(15):8349–54. Epub 2014/05/16. doi: 10.1128/JVI.00924-14. PubMed PMID: 24829350;

58. PubMed Central PMCID: PMC4135926.

58. Horowitz A, Hafalla JC, King E, Lusingu J, Dekker D, Leach A, et al. Antigen-specific IL-2 secretion correlates with NK cell responses after immunization of Tanzanian children with the RTS,S/AS01 malaria vaccine. Journal of immunology. 2012;188(10):5054-62. Epub 2012/04/17. doi: 10.4049/jimmunol.1102710. PubMed PMID: 22504653; PubMed Central PMCID: PMC3378032.

59. Horowitz A, Behrens RH, Okell L, Fooks AR, Riley EM. NK cells as effectors of acquired immune responses: effector CD4+ T cell-dependent activation of NK cells following vaccination. Journal of immunology. 2010;185(5):2808–18. Epub 2010/08/04. doi: 10.4049/jimmunol.1000844. PubMed PMID: 20679529.

60. Major EO, Miller AE, Mourrain P, Traub RG, de Widt E, Sever J. Establishment of a line of human fetal glial cells that supports JC virus multiplication. Proc Natl Acad Sci U S A. 1985;82(4):1257–61. Epub 1985/02/01. doi: 10.1073/pnas.82.4.1257. PubMed PMID: 2983332; PubMed Central PMCID: PMCPMC397234.

61. Vacante DA, Traub R, Major EO. Extension of JC virus host range to monkey cells by insertion of a simian virus 40 enhancer into the JC virus regulatory region. Virology. 1989;170(2):353–61. Epub 1989/06/01. doi: 10.1016/0042-6822(89)90425-x. PubMed PMID: 2543122.

62. Kyrysyuk O, Wucherpfennig KW. Designing Cancer Immunotherapies That Engage T Cells and NK Cells. Annu Rev Immunol. 2022. Epub 2022/11/30. doi: 10.1146/annurev-immunol-101921-044122. PubMed PMID: 36446137.

63. Assetta B, De Cecco M, O’Hara B, Atwood WJ. JC Polyomavirus Infection of Primary Human Renal Epithelial Cells Is Controlled by a Type I IFN-Induced Response. mBio. 2016;7(4). Epub 2016/07/07. doi: 10.1128/mBio.00903-16. PubMed PMID: 27381292; PubMed Central PMCID: PMCPMC4958256.

64. Nolting A, Dugast AS, Rihn S, Luteijn R, Carrington MF, Kane K, et al. MHC class I chain-related protein A shedding in chronic HIV-1 infection is associated with profound NK cell dysfunction. Virology. 2010;406(1):12–20. Epub 2010/07/30. doi: 10.1016/j.virol.2010.05.014. PubMed PMID: 20667578; PubMed Central PMCID: PMC2932841.

65. Matusali G, Tchidjou HK, Pontrelli G, Bernardi S, D’Ettorre G, Vullo V, et al. Soluble ligands for the NKG2D receptor are released during HIV-1 infection and impair NKG2D expression and cytotoxicity of NK cells. FASEB J. 2013;27(6):2440–50. Epub 2013/02/12. doi: 10.1096/fj.12-223057. PubMed PMID: 23395909.

66. Bae DS, Hwang YK, Lee JK. Importance of NKG2D-NKG2D ligands interaction for cytolytic activity of natural killer cell. Cell Immunol. 2012;276(1-2):122–7. Epub 2012/05/23. doi: 10.1016/j.cellimm.2012.04.011. PubMed PMID: 22613008.

67. Sospedra M, Schippling S, Yousef S, Jelcic I, Bofill-Mas S, Planas R, et al. Treating progressive multifocal leukoencephalopathy with interleukin 7 and vaccination with JC virus capsid protein VP1. Clin Infect Dis. 2014;59(11):1588–92. Epub 2014/09/13. doi: 10.1093/cid/ciu682. PubMed PMID: 25214510; PubMed Central PMCID: PMCPMC4650775.

68. Patel A, Patel J, Ikwuagwu J. A case of progressive multifocal leukoencephalopathy and idiopathic CD4+ lymphocytopenia. J Antimicrob Chemother. 2010;65(12):2697–8. Epub 2010/09/25. doi: 10.1093/jac/dkq359. PubMed PMID: 20864499.

69. Miskin DP, Chalkias SG, Dang X, Bord E, Batson S, Koralnik IJ. Interleukin-7 treatment of PML in a patient with idiopathic lymphocytopenia. Neurol Neuroimmunol Neuroinflamm. 2016;3(2):e213. Epub 2016/05/05. doi: 10.1212/NXI.0000000000000213. PubMed PMID: 27144212; PubMed Central PMCID: PMCPMC4841501.

70. Guille M, Rousset S, Bonneville F, Mengelle C, Taoufik Y, Delobel P, et al. IL-7 immunotherapy for progressive multifocal leukoencephalopathy in a patient with already controlled HIV-1 infection on antiretroviral therapy. AIDS. 2019;33(12):1954–6. Epub 2019/09/07. doi: 10.1097/QAD.0000000000002278. PubMed PMID: 31490218.

71. Gasnault J, de Goer de Herve MG, Michot JM, Hendel-Chavez H, Seta V, Mazet AA, et al. Efficacy of recombinant human interleukin 7 in a patient with severe lymphopenia-related progressive multifocal leukoencephalopathy. Open Forum Infect Dis. 2014;1(2):ofu074. Epub 2015/03/04. doi: 10.1093/ofid/ofu074. PubMed PMID: 25734144; PubMed Central PMCID: PMCPMC4281783.

72. Harel A, Horng S, Gustafson T, Ramineni A, Farber RS, Fabian M. Successful treatment of progressive multifocal leukoencephalopathy with recombinant interleukin-7 and maraviroc in a patient with idiopathic CD4 lymphocytopenia. J Neurovirol. 2018;24(5):652–5. Epub 2018/07/11. doi: 10.1007/s13365-018-0657-x. PubMed PMID: 29987583.

73. Nkolola JP, Peng H, Settembre EC, Freeman M, Grandpre LE, Devoy C, et al. Breadth of neutralizing antibodies elicited by stable, homogeneous clade A and clade C HIV-1 gp140 envelope trimers in guinea pigs. J Virol. 2010;84(7):3270–9. Epub 2010/01/08. doi: 10.1128/JVI.02252-09. PubMed PMID: 20053749; PubMed Central PMCID: PMCPMC2838122.

